# Genomic editors for localised population suppression

**DOI:** 10.1101/2025.11.03.686305

**Authors:** Katie Willis, Austin Burt

## Abstract

Effective control of pest populations remains a major challenge for agriculture, public health, and conservation. While genetic control strategies such as the release of sterile males or individuals carrying dominant lethal alleles have achieved some success, they typically require repeated, large-scale releases due to immediate selection against costly alleles, limiting scalability and applicability to species that are difficult to rear in the laboratory. Recent CRISPR/Cas-based genome editing has enabled the development of more efficient genetic control methods that bias their own inheritance and impose a genetic load to achieve population suppression. These designs can potentially spread throughout an entire species range, making them unsuitable when only local or temporary suppression is needed. Here we explore self-limiting genetic strategies based on releasing males carrying autosomal genomic editors that create deleterious edits in essential genes. Unlike traditional approaches, where costly alleles are immediately purged, these editors can persist through multiple generations when creating female-specific or recessive edits, or when editing rates are less than 100%. This allows editors to survive in males, heterozygous carriers, or individuals unaffected by incomplete editing, trading immediate lethality for accumulated genetic load over time. Using deterministic population modelling, we demonstrate that homozygous releases of editors creating female-specific dominant or recessive edits require 45% and 50% fewer males than releasing individuals carrying dominant lethal alleles (RIDL), respectively, to achieve comparable suppression levels. Efficiency gains are further enhanced by targeting multiple genes simultaneously, with editors making female-specific recessive edits showing approximately 50% reduction in release requirements when targeting three genes compared to one – representing a 73% reduction compared to RIDL. Co-releasing editors with a second construct that temporarily boosts editor frequency can achieve efficiency comparable to previously proposed selectively neutral designs while maintaining temporal self-limitation. These results highlight promising alternative strategies for achieving localised, efficient, and self-limiting pest population control suitable for contexts requiring contained suppression.

## INTRODUCTION

Control of pest populations is essential to prevent the spread of vector-borne diseases, crop damage and unwanted environmental change. Genetic biocontrol represents one promising approach, with the sterile insect technique (SIT) being the most widely used to date (Dyck et al., 2021). This involves inundating wild populations with sterile males of the same species that mate with wild females but produce no viable offspring, thereby reducing the population’s reproductive capacity. Traditionally sterile males have been produced through irradiation or chemosterilization, however, advances in molecular techniques have enabled more precise genetic modifications that can induce sterility (Dyck et al., 2021). Examples include RIDL (Release of Insects carrying Dominant Lethals), where males carry genetic constructs that cause offspring lethality; female-specific RIDL (fsRIDL), which targets only female progeny; and precision guided SIT which combines genetic sterility and sex sorting (Alphey, 2002, Kandul et al., 2019). Alternatively, male sterilization has been achieved through *Wolbachia* cytoplasmic incompatibility (IIT), where infected males produce inviable offspring when mating with uninfected females (Mohanty et al., 2016, Crawford et al., 2020, Ching, 2021). While some of these approaches have been successful in several cases (Vreysen et al., 2000, Wyss, 2000, Hendrichs et al., 2002), they can require relatively large and repeated releases to achieve population elimination. This requirement may prove prohibitive for controlling large populations, those geographically hard to reach, or species that are difficult to rear under laboratory conditions.

One way to increase efficiency is to utilise selfish genetic elements to spread costly alleles despite their negative fitness consequence (Burt, 2014, Grilli et al., 2021, Hay et al., 2021, Raban et al., 2023). Two main categories of selfish genetic elements have been proposed for achieving population suppression. First, homing-based constructs that induce a double-strand break at the wildtype version of the construct insertion site, prompting the cells homology directed repair machinery to use the construct as a repair template (process known as homing). This process leads to the construct being copied over, creating biased inheritance if homing occurs in the germline (Burt, 2003). Second, Cleave and Rescue systems involve constructs containing both a genomic editor that creates lethal or sterilizing edits in an essential gene and a recoded copy of the targeted gene, functioning as a toxin-antidote gene drive (Oberhofer et al., 2019, Champer et al., 2020b). When either construct is associated with significant fitness costs – for example, through insertion into and disruption of essential genes – the biased inheritance can be sufficient to outweigh selection against the fitness costs, allowing the costly allele to spread and ultimately result in population suppression (Burt, 2003, Champer et al., 2020a, Simoni et al., 2020, Hammond et al., 2016, Kyrou et al., 2018). For both approaches, only relatively small releases are required because the constructs can increase in frequency from rarity and spread between populations wherever gene flow exists, potentially allowing spread across an entire species range. While species-wide spread is advantageous in some contexts, there is growing interest in identifying strategies which can remain geographically localised while still requiring fewer releases than traditional sterile insect techniques for targeted pest population suppression in specific regions.

One reason for high release requirements of SIT, IIT and RIDL strategies is that the agent responsible for offspring lethality only lasts a single generation. This is equivalent to releasing males homozygous for dominant lethal mutations that lead to immediate death of all progeny. Since the lethal edits are immediately selected against and disappear rapidly, repeated releases are required to achieve and sustain substantial population suppression. Consider instead releasing males carrying a genomic editor that creates costly edits in the germline at target sites elsewhere in the genome. If not all of the released male’s progeny die – whether due to the edits being female-specific, or recessive, or a sub-100% editing efficiency – selection against the editor becomes weaker than selection against a dominant lethal mutation, allowing the editor to persist longer and potentially create additional edits leading to a greater population suppression. Since the editors are inherited in a Mendelian fashion, they are not expected to increase in frequency beyond their initial release ratios or spread extensively beyond the release area. The previously proposed Y-linked editor (YLE; Burt and Deredec, 2018) represents an extreme case, where the editing construct can be completely shielded from selection by being located on the Y chromosome and making female-specific lethal or sterile edits. Protected dominant negative editors (PDNEs) similarly achieve selective neutrality through a single autosomal editor that creates lethal or sterile edits while incorporating a toxin-antidote driving mechanism to counter the negative selection (Willis and Burt, 2025).

In this paper we investigate genetically simpler designs in which the editors and their targets are autosomal, and there is no counter-balancing drive mechanism, and so the editors experience a moderate negative selection pressure due to their association with the edits they create and will decline in frequency over time (but not as rapidly as RIDL constructs). Similar autosomal editors have been proposed for spreading novel traits for population modification or disrupting feminizing genes to convert females to fertile males for population suppression (Johnson et al., 2024). We use mathematical modelling to investigate the predicted efficiency of autosomal editors which disrupt genes essential for survival or fertility, and explore approaches to maximise their effectiveness, by varying the fitness costs of the edits, the number of target sites, the release regime, and pairing the editors with a secondary construct that temporarily boosts editor frequency. Following from previous proposals, these editors could be constructed using CRISPR-based technologies that induce double-strand breaks repaired through non-homologous end joining (Oberhofer et al., 2019, Oberhofer et al., 2021, Metzloff et al., 2021, Oberhofer et al., 2024, Liu et al., 2024), or through base or prime editing that make precise nucleotide modifications (Bosch et al., 2021, Doll et al., 2023, Thakkar et al., 2023, Clark et al., 2024), creating loss-of-function alleles through disruptive insertions and deletions, single nucleotide polymorphisms, or premature stop codons in genes required for survival or fertility.

## RESULTS

### Single releases

We start by considering a single release of editors in homozygous males into a well-mixed population at 100% of the initial male population size, examining four types of editors based on the fitness effects of the edits they make: dominant versus recessive, and affecting both sexes or only females. Edits are assumed to be made at target sites that are unlinked to the construct’s insertion site and cause lethality after the juvenile stage at which density-dependent population regulation occurs, but before reproduction starts, whilst the editor itself is assumed to be inserted into a neutral site.

Figure 1a shows the time course for editor allele frequency, assuming 100% editing efficiency. Editors that create recessive edits (green and magenta) persist for longer than those creating dominant edits (blue and purple), and within each category, editors that create female-specific edits (magenta and purple) persist longer than those that create edits affecting both sexes (blue and green). These differences in persistence occur because the editors can survive into the next generation through males when edits are female-specific, or through heterozygotes when edits are recessive, but are gradually lost when inherited by females or in homozygotes, eventually leading to their disappearance. The same pattern is observed for edit frequency (Figure 1b), with recessive edits initially increasing over successive generations even as the editor itself declines. For comparison, we show the results of releasing males homozygous for the edits rather than the editor, equivalent to RIDL (Figure 1, black) and fsRIDL (Figure 1, grey) approaches. The frequency of an editor making dominant edits affecting both

**Figure 1.**
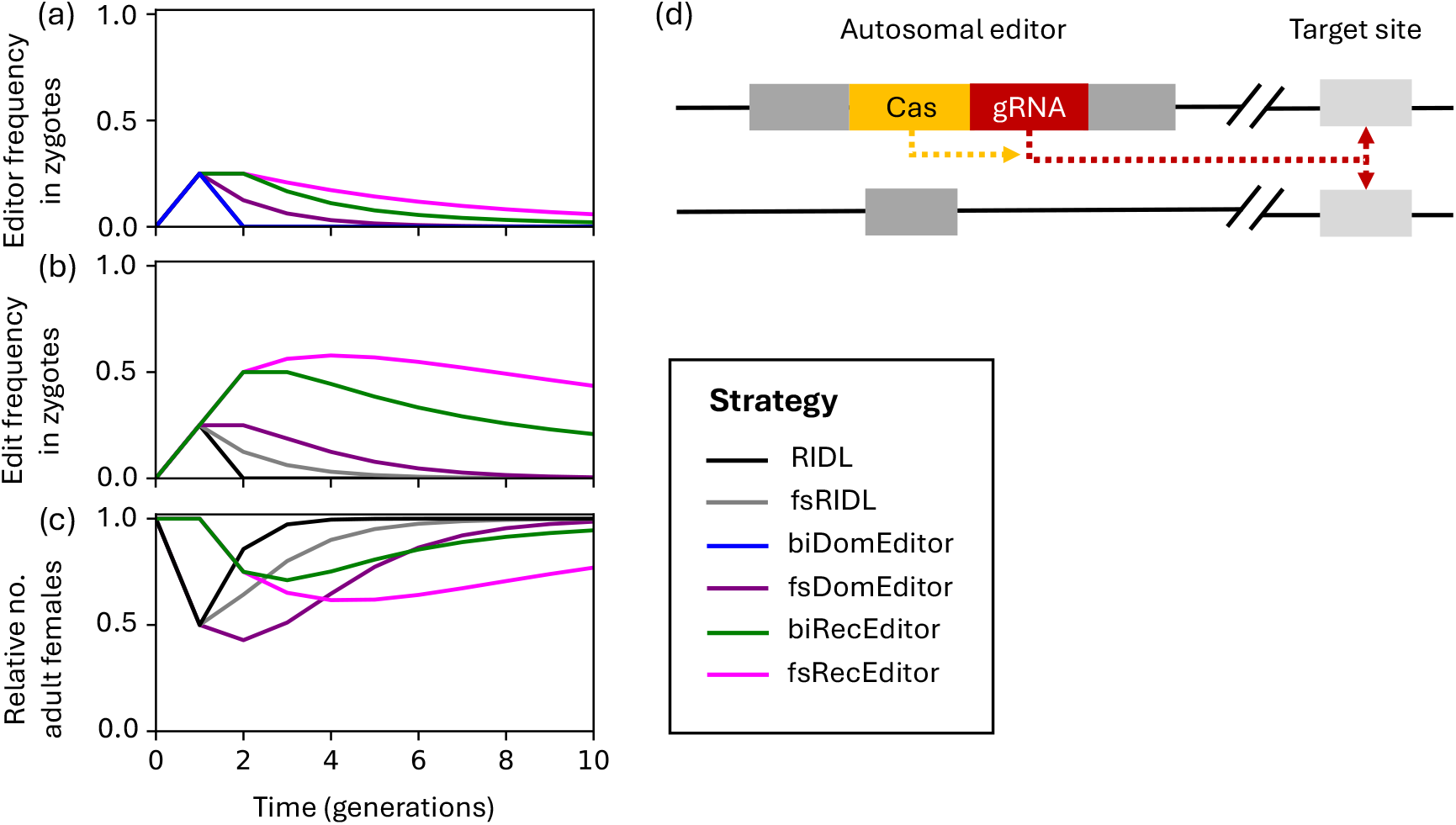
A time series of a single release of a genomic editor that creates edits elsewhere in the genome that cause dominant lethality in both sexes (blue) or only females (purple) or recessive lethality in both sexes (green) or only females (magenta). Shown for comparison are releases of males homozygous for an allele causing dominant lethality in offspring of both sexes (black, equivalent to RIDL) or only females (grey, equivalent to fsRIDL). Males homozygous for the editor (or edit) are released in the first generation at 100% of the initial male population size. **(a)** shows the frequency of the editor in zygotes, **(b)** the frequency of the edit in zygotes and **(c)** the relative number of adult females. Note that the edit frequency and suppression dynamics of editors that create edits causing dominant lethality in both sexes are no different than releasing the edits directly (black lines), since the editor is purged from the population after a single generation. The editing rate of the editor is assumed to be 100% and all fitness costs are maximised. Lethality is assumed to occur after density dependent stages, and the population has an R_m_ of C. Males are released with wildtype versions of the target site, and the editor and its target locus are unlinked. **(d)** shows a possible molecular configuration, where the editor is inserted into a neutral autosomal site and the editors target site is elsewhere in the genome.

sexes, and the frequency of the edits it makes, matches that of a dominant lethal edit when released directly in homozygous males (RIDL, figure 1b, black). This is because with 100% editing efficiency all editors are lost alongside inherited edits and therefore no new edits can be made in subsequent generations. Although female-specific dominant edits released directly (equivalent to fsRIDL) are more persistent than those affecting both sexes, halving in frequency each generation (Figure 1b, grey), they disappear faster than the same edit produced by an editor (Figure 1b, purple). In terms of population size, the female-specific dominant editor achieves the greatest absolute level of suppression (Figure 1c, purple). While the maximum level of suppression achieved by the recessive editors is less, even than RIDL or fsRIDL, the suppression effect is much longer lasting than editors making dominant edits.

### Repeated releases

In each of these designs, repeated releases would typically be required to achieve and sustain effective levels of suppression. One way to compare efficiency under a repeat release regime is to calculate the number of males that need to be released each generation (relative to the starting male population size) to suppress a population by a certain amount. Figure 2 shows the release rates required for a range of designs to suppress a population with an R_m_ of 6 by 95% within 36 generations, assuming 100% editing efficiency. If the edit causes dominant lethality in both sexes (analogous to RIDL), there is no difference between releasing males homozygous for the edit directly versus releasing males heterozygous or homozygous for the editor (Figure 2, blue points, first three panels). However, when the edit causes female-specific dominant lethality, releasing males heterozygous for an editor requires 43% fewer released males than releasing males homozygous for the edit alone (0.40 vs 0.67; Figure 2, first versus second column, purple, one target site). Further gains in efficiency are possible when releasing males homozygous for the editor, requiring only one-third of the release rate compared to releasing the edit alone (0.24 vs 0.67). This strategy also requires 45**%** fewer males to be released than RIDL (0.24 vs 0.44). When the edit is a recessive lethal, whether female-specific or affecting both sexes, releasing males either homozygous for the edit or heterozygous for the editor cannot practically control the population. In contrast, releasing males homozygous for this type of editor achieves efficiency comparable to that observed with editors that make female-specific dominant lethal edits (Figure 2, third column, one target site).

**Figure 2.**
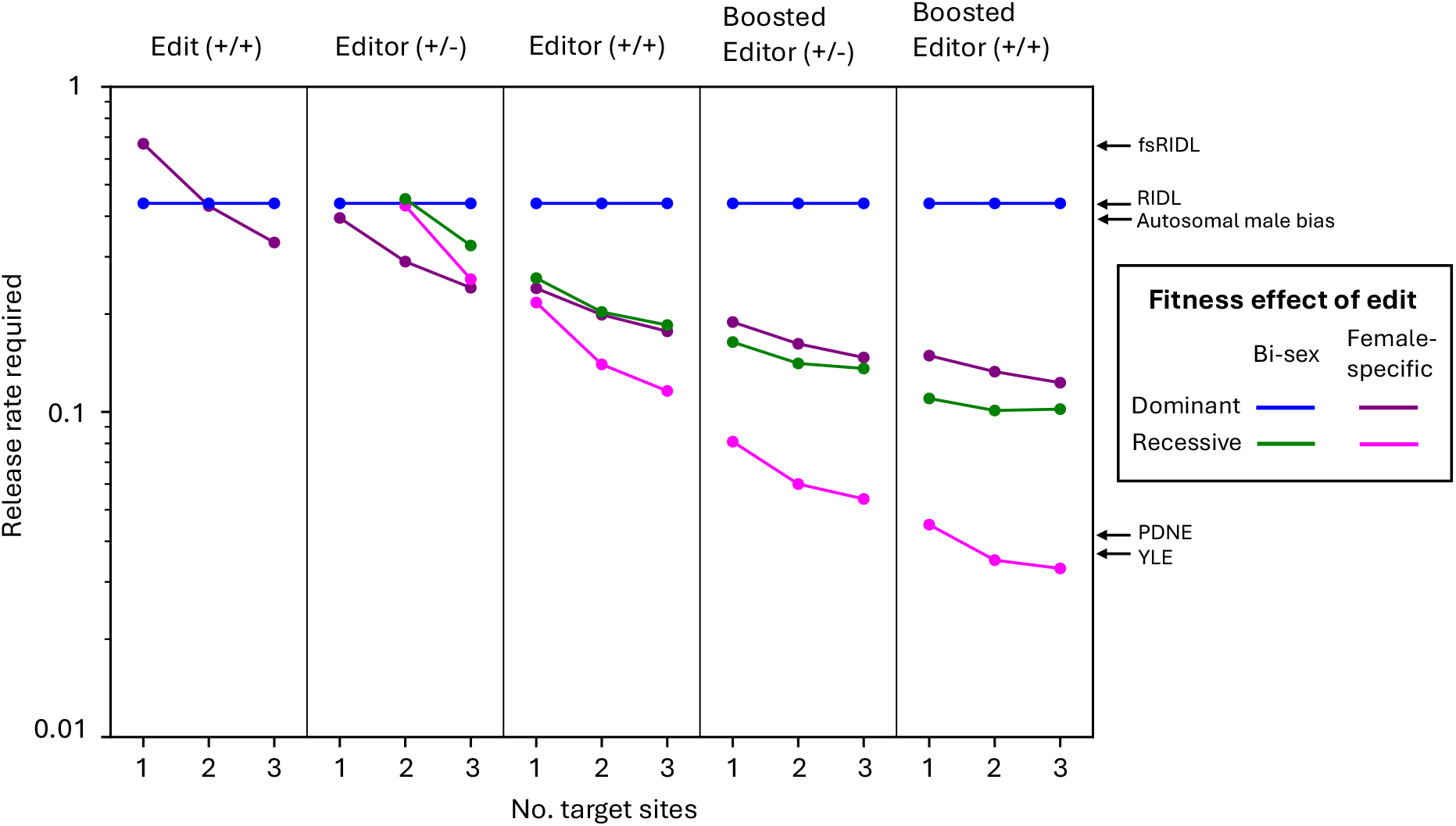
The male release rates required (relative to the starting male population size) to suppress a population with an R_m_ of C by S5% within 3C generations for a range of different strategies. Labelled columns indicate different designs, including releasing males carrying two copies of an edit, one (+/-) or two (+/+) copies of an editor and one or two copies of an editor released alongside a homing-based booster construct. Editor (and edits) can vary in their fitness effects, causing female-specific dominant (purple) or recessive (magenta) lethality, or bi-sex dominant (blue) or recessive (green) lethality. For comparison, arrows on the right indicate release rates for previously proposed designs including fsRIDL, RIDL, a sex-ratio distorter located in an autosome, a protected dominant negative editor (PDNE) and a Y-linked editor (YLE). The following combinations are not shown due to impractical high release rate requitements: homozygous releases of recessive edits and heterozygous releases of an editor making recessive edits at a single target site. Editing is assumed to be at 100% efficiency, all fitness parameters are maximized, and constructs and target sites are unlinked.

### Targeting multiple sites

One way to further increase the efficiency of editors is to design them to make edits in more than one target gene. For CRISPR-based genomic editors, this could be achieved by incorporating multiple gRNAs into the editor construct to target multiple essential genes, each of which independently causes reduced survival or fertility when edited. To evaluate this approach, we modelled scenarios where multiple target genes are unlinked to each other and to the editor. In all designs other than an editor that makes edits causing dominant lethality in both sexes, creating edits in more than one gene always improves efficiency (Figure 2). This increased efficiency stems from a greater total edit burden in the population and independent segregation of edits at multiple loci. Targeting multiple unlinked sites increases the proportion of offspring inheriting at least one deleterious edit compared to single-site targeting, thereby amplifying the genetic load imposed. The strategy showing the most improvement in efficiency due to additional target loci is homozygous releases of an editor making female-specific recessive edits, where targeting more than one gene offers a clear efficiency advantage over all alternative strategies (Figure 2, third column, magenta). Release rate requirements fall by approximately 47% when targeting three genes compared to one (0.12 vs 0.22), compared to an approximate 27% reduction in release rate for editors creating recessive edits affecting both sexes or dominant female-specific edits (0.185 vs 0.26 and 0.18 vs 0.24 respectively). When compared to RIDL, releasing homozygous males carrying an editor creating female-specific recessive edits at two or three genes requires 68**%** (0.14 vs 0.44) and 73% (0.12 vs 0.44) smaller releases respective.

### Releasing with a booster

Another way to improve efficiency is to release the genomic editor with a trans-acting booster construct that allows the editor to home in its presence, enabling the construct to reach greater frequency in the population, persist for longer and consequently create more edits while imposing a greater genetic load. This could be engineered by placing a gRNA in an unlinked locus that targets the editor’s insertion site, such that the editor homes in the presence of the booster. Since the booster is inherited in a Mendelian manner, it is eventually lost over time, making the boosting effect temporary. Our modelling of this system, assuming the booster is unlinked to the editor and causes homing of the editor at 100% efficiency, shows this approach to be most advantageous when applied to an editor that makes female-specific recessive edits, reducing required release rates by 70-80% compared to homozygous release of the editor alone, depending on the number of target sites (TS) (Figure 2, fourth and fifth column; TS = 1, 0.05 vs 0.22; TS=2, 0.04 vs 0.14; TS=3, 0.03 vs 0.12). The greater advantage for female-specific recessive editors is because the booster can persist longest when paired with this design, surviving in males and heterozygotes for the edit, thus enabling the booster to elevate the editor to greater frequency. This strategy brings release rates down to levels comparable to other best-in-class designs for localised suppression such as the YLE and PDNE, despite the editor not being selectively neutral when optimally designed. From a practical standpoint, the boosting action makes heterozygous releases viable for editors creating both types of recessive edit, which may be easier to generate in the laboratory, whilst also reducing release rates by 53-63% compared to homozygous releases of an editor alone (0.16 vs 0.26 for edits affecting both sexes and 0.08 vs 0.22 for those that are female-specific).

### Sensitivity to variation in editing rates

In reality it is unlikely that an editor would create edits 100% of the time as we have modelled so far, therefore we next performed a sensitivity analysis on each strategy, varying the editing rate (Figure 3). For sub-100% editing scenarios, we assume editors are built using base or prime editing systems (and therefore there is no homing of edits made in previous generations), and that failed editing attempts leave the target site intact for potential future editing. In general, all designs are resilient to less-than-perfect editing when the editing rate remains above a “failure threshold”, below which the strategy fails to practically suppress the population by the target 95% within the 36-generations. Homozygous releases, boosting, and increasing the number of target sites all reduce this failure threshold compared to heterozygous releases, with the most resilient designs being effective with approximately 30% editing efficiency. Interestingly, lower editing rates can slightly improve efficiency in cases where the editor is released in homozygotes (Figure 3, lower) or for boosted designs (Figure 3, thick lines). This also occurs for editors that create dominant lethal edits affecting both sexes, which when editing is maximised is no better than releasing the edits directly (Figure 3, blue). This counterintuitive advantage arises from increased persistence of the editor in the population. With lower editing rates, the editor is less frequently inherited alongside an edit it creates, reducing the likelihood that it is purged through selection against the costly edit. This permits the editor to persist for longer in the population, and in some cases results in more total edits being made within the suppression timeframe, ultimately imposing a greater genetic load. When editing is less than perfect, linkage between the editor and its target site(s) reduces efficiency, however this effect is relatively minor (Supplementary Feigure 1).

**Figure 3.**
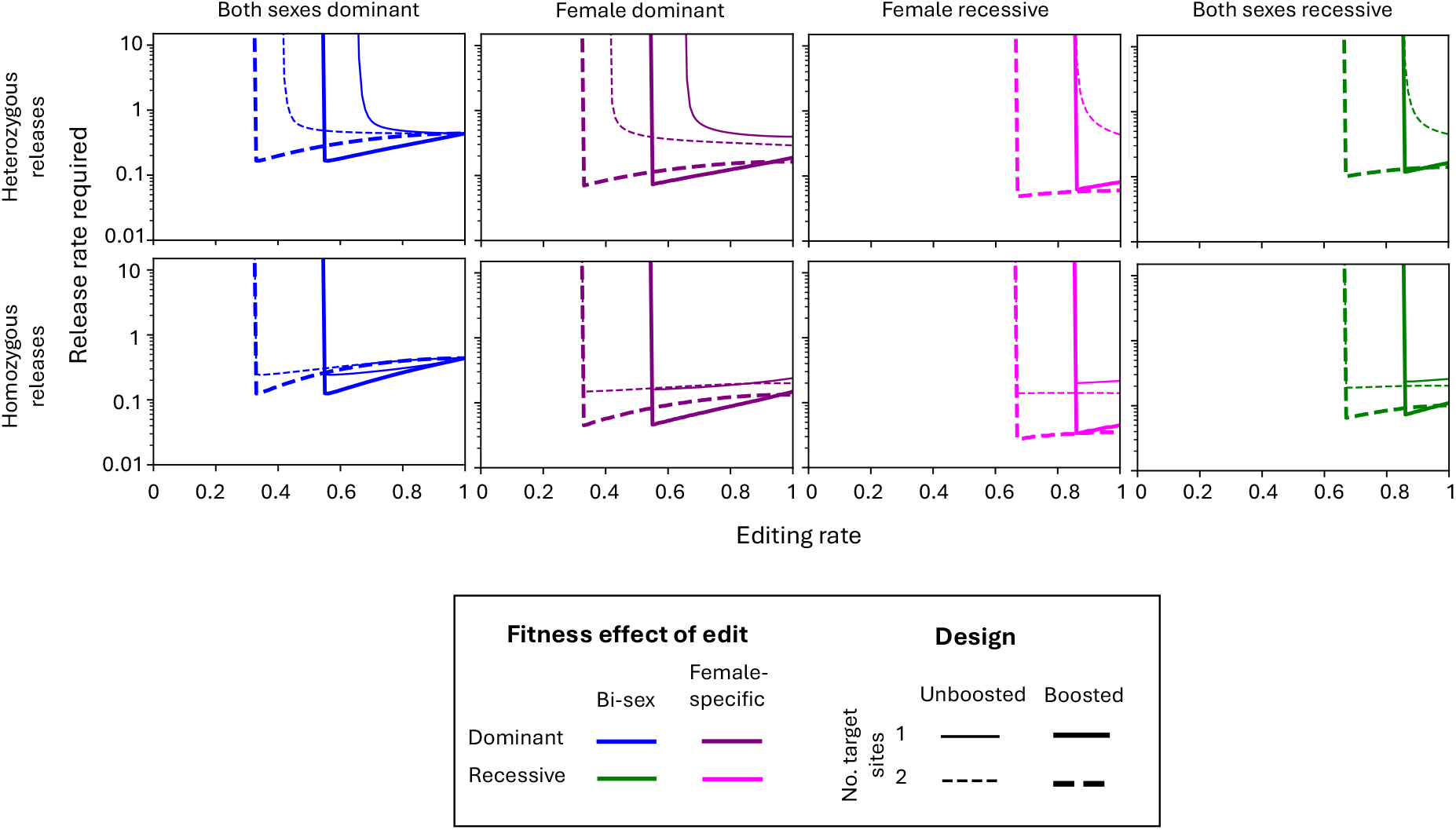
The male release rates required relative to the starting male population size in order to suppress a population with an R_m_ of C by S5% within 3C generations for a range of different autosomal editor strategies whilst varying the rate at which edits are created. Variations of the designs include releasing males heterozygous (top row) or homozygous (bottom) for the editor, those where the edit causes dominant lethality in both sexes (blue), in only females (purple), or recessive lethality in both sexes (green) or only females (magenta). The number of edited genes also vary including one (solid) or two (dashed) genes, and the editor is released alone (thin lines) or with a booster which causes the editor to home (thicker lines). The editing rate per wildtype target site is assumed to be the same whether editing occurs in heterozygotes or homozygotes for the target site. The homing efficiency induced by the booster construct is assumed to be 100% and all fitness parameters are maximized.

### Optimal release regimes

Given that each design has an effect over different time frames (Figure 1), we next investigated the optimal combination of release number and size required for each strategy to achieve a certain level of suppression. For each strategy, we calculated the release rate required (one release each generation) to suppress the population by 95% when released over a range of different periods (2 to 40 generations) and identified the combination of release number and size that minimised the total number of males released (Figure 4). In general, editors that create dominant edits work most efficiently when releasing larger numbers of males over fewer generations (Figure 4, purple; between 0.28 and 1.89 release rate for 5-8 generations), while editors that make recessive edits achieve optimal efficiency when releasing smaller numbers over more generations (Figure 4, pink and green; between 0.18 to 0.81 release rate for 9-12 and between 0.06 to 0.77 release rate for 11-16 generations for edits affecting both sexes or only female respectively). This difference reflects the temporal dynamics of these approaches observed in Figure 1: recessive edits require an additional generation to impact population growth since they only affect homozygous individuals, and the editors creating them tend to persist longer in the population. Consequently, longer release timeframes allow both the editor and the resulting edits to accumulate, leading to greater load imposed. The total number of males required for release over the entire period varied by just over an order of magnitude across all explored designs, ranging from 0.8 to 10 times the starting male population. Female-specific recessive designs span this entire range, while the female-specific dominant designs show a narrower range between 2.2 and 7.7 times the starting population (assuming 100% editing efficiency). When comparing editors released as homozygotes, similar total release numbers are required regardless of fitness effect (female-specific dominant 2.7 – 3.3, female-specific recessive 2.5 - 4.8 and bi-sex recessive 3.3 – 4.8, with ranges reflecting the number of target sites). However, boosted designs of the female-specific recessive editor offer a two-fold reduction in total release requirements compared to the dominant equivalent (female-specific dominant 2.2 – 2.5; female-specific recessive 0.8 – 1.3), aligning with differences in efficiency seen in Figure 2.

**Figure 4.**
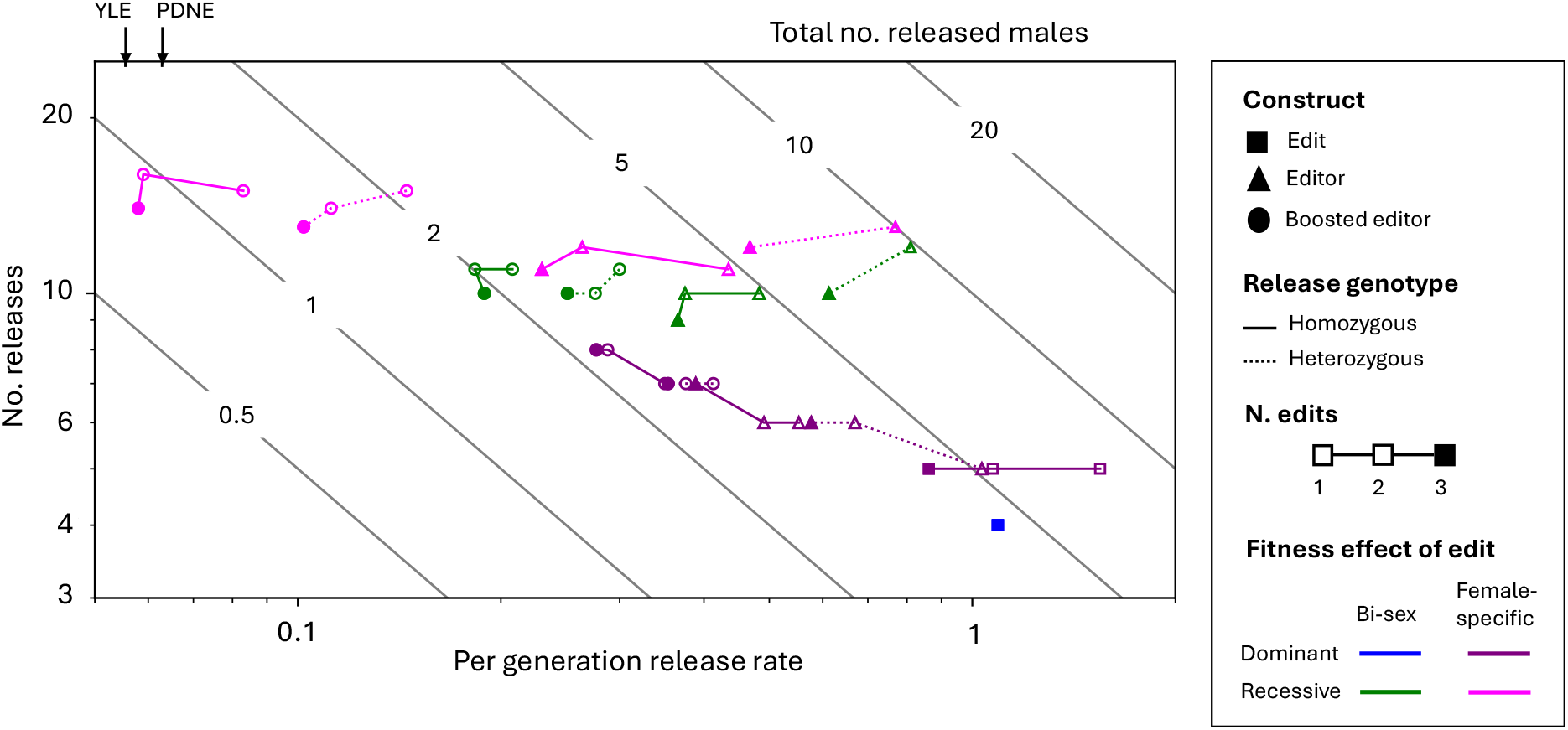
The optimal combination of release size and release number that minimizes the total number of males needed for release to suppress a population with an R_m_ of C by S5% for a range of different designs. Diagonal isoclines (grey) represent different combinations of per generation release rate and number of releases that generate the same total number of males released over the whole release period. Strategies include releasing males homozygotes (solid lines) or heterozygous (dashed lines) for an edit (square), an editor (triangle) or a boosted editor (circle). Black lines link together designs which differ only in their number of target sites, with the filled marker indicating 3 target sites to give direction. The following combinations are not shown due to impractical high release requirements: heterozygous releases of an edit, homozygous releases of a recessive edit and heterozygous releases of an editor making recessive edits at a single target site. For comparison, the per generation release rate for males carrying a YLE or PDNE is shown when released for 25 generations. Since both are selectively neutral, the efficiency will always improve when released over more generations; however, this efficiency gain reduces as numbers of releases increase, with the total number of released males reducing by <0.01 each additional release over 25 generations. All efficiencies and intended fitness costs are at 100% and target sites are assumed unlinked.

## DISCUSSION

Our results demonstrate that releasing males carrying autosomal genomic editors that create lethal mutations can give improved efficiency compared to releasing males carrying lethal mutations directly (equivalent to SIT or RIDL approaches) by extending the persistence of load-inducing constructs across multiple generations. The approach depends on the uncoupling of the editor construct and the costly edits it creates, allowing the editors to persist through individuals that are not affected by the edits they produce. This persistence is achieved when editors create female-specific or recessive edits, allowing the construct to survive in males or heterozygous carriers respectively, or when editing rates are less than 100%. Similar approaches have been taken to increase persistence of released mutations themselves (e.g. in RIDL versus fsRIDL), where female-specific mutations can persist over multiple generations through males (Labbé et al., 2012). For the genomic editors described here, persistence of the costly edits is further enhanced since the construct can create new edits over successive generations, leading to prolonged population suppression, and reduced release requirements. Our findings reveal that homozygous releases of editors creating female-specific dominant or recessive edits require 45% and 50% fewer males than RIDL respectively to achieve comparable suppression levels. For editors targeting multiple sites, this efficiency advantage becomes even more pronounced, particularly for editors creating female-specific recessive edits, showing a 73% reduction compared to RIDL.

Several other genomic editor-based control strategies have been proposed, some specifically for localised population suppression. YLEs and PDNEs induce a reproductive load upon a population while achieving selective neutrality by either placing the editor on the Y chromosome to avoid selection against the female-specific edits it makes, or using a toxin-antidote drive system to balance the negative selection (Burt and Deredec, 2018, Tolosana et al., 2024, Willis and Burt, 2025). When parameters are optimized, these approaches can persist indefinitely at stable frequencies, providing sustained population suppression. By contrast, even under idealised conditions, our autosomal editors decline in frequency over time due to their association with the costly edits they create. While autosomal genomic editors potentially require greater repeated releases due to stronger negative selection, when released alongside a homing-based booster (genetically not-linked to the editor, i.e. non-autonomous), editors creating female-specific recessive edits can achieve comparable suppression levels with similar release rates to YLEs and PDNEs, but are expected to disappear from the population more quickly. The Ifegenia (inherited female elimination by genetically encoded nucleases to interrupt alleles) system similarly involves releasing males carrying constructs expressing Cas9 and gRNA targeting female-specific genes, and has been demonstrated in *Anopheles gambiae*, targeting the essential gene *femaleless* with 85-95% editing efficiency (Smidler et al., 2023). Though similar, this design differs from our autonomous editors as it employs unlinked Cas9 and gRNA components, creating a non-autonomous system that requires both components and a WT target allele to be present for editing to occur. Split drive systems, where gRNA or Cas components are inserted into the essential target gene itself, represent another variant of this approach, as well as those which involve a chain of “booster” construct, each causing the other to drive (Noble et al., 2019, Guo et al., 2025). Further modelling would be required to evaluate how our autonomous designs compare to these alternatives.

Due to the editor being selected against when associated with the edits it makes our analysis revealed a trade-off between immediate population suppression and long-term persistence. Editors creating dominant edits achieve greater immediate suppression but persist for shorter duration, while those creating recessive edits maintain lower peak suppression but extend their effects over longer timeframes. This trade-off impacts optimal release regimes: dominant editors perform best with fewer large releases, while recessive editors are most effective through many smaller releases over longer periods. The trade-off also explains why editors creating female-specific recessive edits are most advantaged by booster pairing. These editors receive the least negative selection of all designs due to the combined recessive and female-specific nature of their edits, minimising selection against the booster and permitting maximal boosting effects. Here, increased longevity compensates for reduced per-generation effectiveness, resulting in greater cumulative genetic load over the suppression timeframe. This persistence advantage similarly explains the counterintuitive observation that editing rates below 100% can improve overall strategy effectiveness. Reduced editing rates decrease the likelihood of the editor being inherited alongside a costly edit, thereby enhancing its persistence and allowing more total edits to accumulate over time. This finding contrasts with previous modelling work on equivalent autosomal genomic editors for population modification or replacement (allele sails), where authors found that low editing rates reduced effectiveness (Johnson et al., 2024). The difference likely stems from the higher fitness costs imposed by our suppression-focused edits and the repeat release regimens employed, where persistence becomes more valuable than immediate efficiency. Johnson et al. (2024) also demonstrated that autosomal editor dynamics are affected by whether editing occurs in the soma or involves maternal deposition of the editing machinery. While not modelled directly, we expect additional costs from somatic editing would increase selection against the editor, reduce persistence, and therefore increase required release numbers. Conversely, maternal deposition would allow creation of additional edits without directly inducing selection against the editor allele, potentially increasing the genetic load with minimal negative consequence for editor persistence. Further modelling would be needed to evaluate how big an affect these processes have on efficiency, and how they differ between different editor designs.

One way to engineer an editor to induce sterility or lethality is to create loss-of-function alleles in haplo-sufficient or -insufficient genes by inserting premature stop codons or indels that cause frameshifts in coding regions. CRISPR-induced double-strand breaks have been used to achieve loss-of-function alleles in the germline with rates of up to 100% in *Drosophila* and *Arabidopsis* (Oberhofer et al., 2019, Oberhofer et al., 2021, Metzloff et al., 2021, Oberhofer et al., 2024, Liu et al., 2024), but sequence changes can be unpredictable (Anzalone et al., 2020). Base editors offer more precise editing, and germline insertion of stop codons has been demonstrated in *Drosophila* with 90%-100% editing rates, causing male-specific sterility or female-specific lethality (Clark et al., 2024), or to induce loss-of-phenotype in non-essential genes (Doll et al., 2023, Thakkar et al., 2023). Prime editors provide similar precision but currently with lower efficiencies in *Drosophila* (∼37%) (Bosch et al., 2021).

Though precise, both approaches can create unintended nucleotide substitutions and indels (Anzalone et al., 2020, Doll et al., 2023). If a fully functional resistant allele arose, we expect it would be strongly selected for and spread to fixation, preventing further editing. Once the wild population is fixed for the resistant allele, releases could still cause suppression, but would work much like releasing the edits directly, since the only WT alleles available for editing would be those present in the released males.

Nevertheless, if significant suppression occurred during spread of resistance, releases into a resistant but substantially smaller population could be enough to maintain suppression using many fewer males than required when only releasing edits directly. Overall efficiency would still be reduced compared to cases where functional resistance did not arise, and the probability of avoiding it could be increased by selecting highly conserved target sites, targeting multiple sites within each target gene or by choice of Cas protein (Kyrou et al., 2018, Oberhofer et al., 2019, Champer et al., 2020c, Yang et al., 2022, Hillary and Ceasar, 2023, Chen et al., 2024, Morianou et al., 2024). Editor designs targeting multiple genes may, in addition to increased efficiency, further reduce probability of resistance spreading, and the impact if it does spread. If edits with alternative fitness costs than those intended are made, such as dominant rather than recessive, or bisex rather than female-specific, the efficiency of the strategy is likely to fall in between each idealised case, though more modelling would be needed to explore this further.

Previously there have been proposals for combining load-inducing editors with autonomous drive systems, to facilitate spread, make suppression more efficient, or even to be the primary source of load. These include proposals to add load-inducing editors to driving Y chromosomes (Deredec et al., 2011), the t-haplotype in mice (Gierus et al., 2022), and a homing suppression drive (Faber et al., 2024). Combining the autosomal editor described in this paper with an autonomous driver would lead to significantly reduced release requirements but could spread wherever there is ongoing gene flow. A geographically or sub-species restricted driving editor could be achieved by using a CRISPR-based drive, where the editor construct contains an additional gRNA targeting its own insertion site which is also locally enriched in the target population (Sudweeks et al., 2019, Willis and Burt, 2021, Geci et al., 2022). This would permit the editor to drive from low frequency in the target population but to remain rare in the non-target population. Analogous to our previously proposed autosomal double drives (Willis and Burt 2019), separation of the locus that the construct homes into and the locus where load is induced would reduce selection against the editor construct, making it more resilient in the face of resistance. Compared to a double drive, an editor-based design would involve only a single genetic construct, creating small edits elsewhere in the genome.

In summary, autosomal genomic editors represent a promising intermediate between traditional sterile male releases and autonomous gene drives. Due to their multi-generation effect, these editors achieve enhanced efficiency compared to SIT, IIT and RIDL while remaining self-limiting and localised. Their robustness to imperfect editing and compatibility with multiple molecular platforms make them potentially applicable for a range of pest species, though the choice of specific design will depend on the specific target pest population and timeframe within which the suppression is required. Though our modelling has identified optimal time frames to minimise release numbers, it does not consider economic costing of rearing and release of small versus large numbers of males, release regimes that skip generations, or the geographical distribution of release sites. These factors may alter the optimal release regimes for each genetic design, and more bespoke, case-by-case modelling would be required to identify which strategy would be most suited.

## METHODS

To simulate the impact of releasing the construct(s) into a wild population, we developed a series of deterministic models which simulate a single, panmictic population of infinite size with discrete, non-overlapping generations, based on that modelled by Burt and Deredec (2018). We began with a baseline model involving two autosomal loci, both with two alleles. At the first locus the first allele is a wildtype allele and the second a genomic editor. At the second locus the first allele is a wildtype allele, susceptible to editing by the genomic editor, and the second is the edited allele. Males and females are modelled separately, resulting in 10 genotypes of each sex. In genotypes that have at least one editor and one wildtype target allele, the editor converts each wildtype target allele with probability *μ*, converting the wildtype allele to an edited allele. We assume no difference in editing rate between individuals with one or two copies of the editor. The fitness effects of the edited allele are modelled by parameters *S*_*X*_ and *h*_*X*_, where *S*_*X*_ describes the fitness cost of the edit in homozygotes and *h*_*X*_ is the dominance coefficient describing the proportion of homozygous costs that affect heterozygotes, both of which can differ between sexes (*X*). The editor itself is assumed to be neutral with no impact on fitness. We also modelled release of the edits alone, varying the fitness parameters and setting the editing rate to zero (*μ =* 0).

To model multiple target loci, the baseline model was extended to include 2 or 3 target loci, each with two alleles (wildtype allele susceptible to editing and edited allele), resulting in a total of 36 and 136 genotypes per sex respectively. In the presence of the editor, each wildtype allele at each target locus has a probability *μ* of being converted to an edited allele. To model linkage between loci, all loci are assumed to be arranged linearly along a single chromosome, where recombination between each pair of loci can occur with probability *r*_*ij*_, where *i* and *j* represent two adjacent loci. To model a pair of loci on different chromosomes *r*_*ij*_ *=* 0.5, whereas when *r*_*ij*_ *=* 0 loci are tightly linked.

To model the editor when paired with a homing-based booster an additional locus was included in each model, containing two alleles: the wildtype and booster construct.

This resulted in a total of 36, 136 and 528 genotypes per sex for editors with one, two and three target sites respectively. In heterozygotes for the editor and a wildtype allele at the same locus, and in the presence of a one or more copies of the booster, homing of the editor occurs with probability *c*, converting the wildtype allele to an editor, where *c =* 1 in all cases modelled. For all results the booster was assumed to be autosomal, unlinked from both the editor and target site(s) and non-autonomous, meaning that the booster cannot cleave its target site in the absence of the editor construct.

To evaluate the impact on population size, we include a population dynamics model which monitors the number of males and females overtime, tracking larvae, pupae and adults. In each generation, females produce *f* eggs. These are fertilised by males, assuming that the number of males does not limit fertilisation success and that all mating is random. Density dependent mortality is applied at the larvae to pupae transition, with a survival probability of 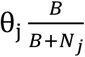 according to the Beverton-Holt model, where θ_*j*_ is the density independent probability of surviving the larval stage, *N*_*j*_ is the total number of larvae and *B* determines the strength of density dependent mortality. All results are reported as population size relative to the starting population, and therefore the value of *B* does not matter. The intrinsic rate of increase is set to *R*_*m*_ *=* 6, and equivalent to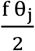. All genotype-dependent fitness costs were applied after density dependent effects, at the pupae to adult transition, assuming pupae die before they emerge as adults. All releases were of adult males, either heterozygous or homozygous for the editor, booster, edit or combinations. Where multiple constructs were released, i.e. the editor and booster, they were released in the same males. All allele frequencies were reported at the zygote (larval) stage, calculated in males and females independently and then averaged. All simulations were performed in Julia, a scientific programming language within the Jupyter notebook interface.

## Notes

### Competing Interest Statement

The authors have declared no competing interest.

### Funding

- Gates Foundation
- Open Philanthropy Project

## REFERENCES

Alphey, L. 2002. Re-engineering the sterile insect technique. Insect Biochem Mol Biol, 32, 1243–7.

Anzalone, A. V., Koblan, L. W. C Liu, D. R. 2020. Genome editing with CRISPR–Cas nucleases, base editors, transposases and prime editors. Nat Biotech, 38, 824–844.

Ashburner, M., Golic, K. G. C Hawley, R. S. 2005. Drosophila : a laboratory handbook, Cold Spring Harbor, N.Y., Cold Spring Harbor Laboratory Press.

Bosch, J. A., Birchak, G. C Perrimon, N. 2021. Precise genome engineering in Drosophila using prime editing. Proc Natl Acad Sci, 118, e2021996118.

Burt, A. 2003. Site-specific selfish genes as tools for the control and genetic engineering of natural populations. Proc Roy Soc Lond B, 270, 921–928.

Burt, A. 2014. Heritable strategies for controlling insect vectors of disease. Philos Trans R Soc Lond B, 369.

Burt, A. C Deredec, A. 2018. Self-limiting population genetic control with sex-linked genome editors. Proc Roy Soc Lond B, 285.

Champer, J., Kim, I. K., Champer, S. E., Clark, A. G. C Messer, P. W. 2020a. Performance analysis of novel toxin-antidote CRISPR gene drive systems. BMC Biol, 18, 27.

Champer, J., Lee, E., Yang, E., Liu, C., Clark, A. G. C Messer, P. W. 2020b. A toxin-antidote CRISPR gene drive system for regional population modification. Nat Commun, 11.

Champer, J., Yang, E., Lee, E., Liu, J. X., Clark, A. G. C Messer, P. W. 2020c. A CRISPR homing gene drive targeting a haplolethal gene removes resistance alleles and successfully spreads through a cage population. Proc Natl Acad Sci, 117, 24377–24383.

Chen, W., Guo, J., Liu, Y. C Champer, J. 2024. Population suppression by release of insects carrying a dominant sterile homing gene drive targeting doublesex in Drosophila. Nat Commun, 15, 8053.

Ching, N. L. 2021. Wolbachia-mediated sterility suppresses Aedes aegypti populations in the urban tropics. medRxiv, 2021.06.16.21257922.

Clark, M., Nguyen, C., Nguyen, H., Tay, A., Beach, S. J., Maselko, M. C López Del Amo, V. 2024. Expanding the CRISPR base editing toolbox in Drosophila melanogaster. Commun Biol, 7, 1126.

Crawford, J. E., Clarke, D. W., Criswell, V., Desnoyer, M., Cornel, D., Deegan, B., Gong, K., Hopkins, K. C., Howell, P., Hyde, J. S., Livni, J., Behling, C., Benza, R., Chen, W., Dobson, K. L., Eldershaw, C., Greeley, D., Han, Y., Hughes, B., Kakani, E., Karbowski, J., Kitchell, A., Lee, E., Lin, T., Liu, J., Lozano, M., Macdonald, W., Mains, J. W., Metlitz, M., Mitchell, S. N., Moore, D., Ohm, J. R., Parkes, K., Porshnikoff, A., Robuck, C., Sheridan, M., Sobecki, R., Smith, P., Stevenson, J., Sullivan, J., Wasson, B., Weakley, A. M., Wilhelm, M., Won, J., Yasunaga, A., Chan, W. C., Holeman, J., Snoad, N., Upson, L., Zha, T., Dobson, S. L., Mulligan, F. S., Massaro, P. C White, B. J. 2020. Efficient production of male Wolbachia-infected Aedes aegypti mosquitoes enables large-scale suppression of wild populations. Nat Biotechnol, 38, 482–492.

Deredec, A., Godfray, H. C. J. C Burt, A. 2011. Requirements for effective malaria control with homing endonuclease genes. Proc Natl Acad Sci, 108, E874–E880.

Doll, R. M., Boutros, M. C Port, F. 2023. A temperature-tolerant CRISPR base editor mediates highly efficient and precise gene editing in Drosophila. Science Advances, 9, eadj1568.

Dyck, V. A., Hendrichs, J. C Robinson, A. S. 2021. Sterile Insect Technique : Principles And Practice In Area-Wide Integrated Pest Management, Taylor and Francis.

Faber, N. R., Xu, X., Chen, J., Hou, S., Du, J., Pannebakker, B. A., Zwaan, B. J., Van Den Heuvel, J. C Champer, J. 2024. Improving the suppressive power of homing gene drive by co-targeting a distant-site female fertility gene. Nat Commun, 15, 9249.

Geci, R., Willis, K. C Burt, A. 2022. Gene drive designs for efficient and localisable population suppression using Y-linked editors. PLoS Genetic, 18, e1010550.

Gierus, L., Birand, A., Bunting, M. D., Godahewa, G. I., Piltz, S. G., Oh, K. P., Piaggio, A. J., Threadgill, D. W., Godwin, J., Edwards, O., Cassey, P., Ross, J. V., Prowse, T. A. A. C Thomas, P. Q. 2022. Leveraging a natural murine meiotic drive to suppress invasive populations. Proc Natl Acad Sci, 119, e2213308119.

Grilli, S., Galizi, R. C Taxiarchi, C. 2021. Genetic technologies for sustainable management of insect pests and disease vectors. Sustainability, 13.

Guo, J., Chen, W. C Champer, J. 2025. Experimental demonstration of daisy chain gene drive and modelling of daisy suppression systems. bioRxiv, 2025.09.20.677490.

Hammond, A., Galizi, R., Kyrou, K., Simoni, A., Siniscalchi, C., Katsanos, D., Gribble, M., Baker, D., Marois, E., Russell, S., Burt, A., Windbichler, N., Crisanti, A. C Nolan, T. 2016. A CRISPR-Cas9 gene drive system-targeting female reproduction in the malaria mosquito vector Anopheles gambiae. Nat Biotechnol, 34, 78–83.

Hay, B. A., Oberhofer, G. C Guo, M. 2021. Engineering the composition and fate of wild populations with gene drive. Annu Rev Entomol, 66, 407–434.

Hendrichs, J., Robinson, A. S., Cayol, J. P. C Enkerlin, W. 2002. Medfly areawide sterile insect technique programmes for prevention, suppression or eradication: the importance of mating behaviour studies. Fla Entomol, 85, 1-13, 13.

Hillary, V. E. C Ceasar, S. A. 2023. A Review on the Mechanism and Applications of CRISPR/Cas9/Cas12/Cas13/Cas14 Proteins Utilized for Genome Engineering. Mol Biotechnol, 65, 311–325.

Johnson, M. L., Hay, B. A. C Maselko, M. 2024. Altering traits and fates of wild populations with Mendelian DNA sequence modifying Allele Sails. Nat Commun, 15, 6665.

Kandul, N. P., Liu, J., Sanchez C H. M., Wu, S. L., Marshall, J. M. C Akbari, O. S. 2019. Transforming insect population control with precision guided sterile males with demonstration in flies. Nat Commun, 10, 84.

Kyrou, K., Hammond, A. M., Galizi, R., Kranjc, N., Burt, A., Beaghton, A. K., Nolan, T. C Crisanti, A. 2018. A CRISPR-Cas9 gene drive targeting doublesex causes complete population suppression in caged Anopheles gambiae mosquitoes. Nat Biotechnol, 36, 1062.

Labbé, G. M. C., Scaife, S., Morgan, S. A., Curtis, Z. H. C Alphey, L. 2012. Femalespecific flightless (fsRIDL) phenotype for control of Aedes albopictus. PLoS Negl Trop Dis, 6, e1724.

Liu, Y., Jiao, B., Champer, J. C Qian, W. 2024. Overriding Mendelian inheritance in Arabidopsis with a CRISPR toxin-antidote gene drive that impairs pollen germination. Nat Plants, 10, 910–922.

Metzloff, M., Yang, E., Dhole, S., Clark, A. G., Messer, P. W. C Champer, J. 2022. Experimental demonstration of tethered gene drive systems for confined population modification or suppression. BMC Biol, 20, 119.

Mohanty, I., Rath, A., Mahapatra, N. C Hazra, R. K. 2016. Wolbachia: A biological control strategy against arboviral diseases. J Vector Borne Dis, 53, 199–207.

Morianou, I., Phillimore, L., Khatri, B. S., Marston, L., Gribble, M., Burt, A., Bernardini, F., Hammond, A. M., Nolan, T. C Crisanti, A. 2024. Engineering Resilient Gene Drives Towards Sustainable Malaria Control: Predicting, Testing and Overcoming Target Site Resistance. bioRxiv, 2024.10.21.618489.

Noble, C., Min, J., Olejarz, J., Buchthal, J., Chavez, A., Smidler, A. L., Debenedictis, E. A., Church, G. M., Nowak, M. A. C Esvelt, K. M. 2019. Daisy-chain gene drives for the alteration of local populations. Proc Natl Acad Sci, 116, 8275–8282.

Oberhofer, G., Ivy, T. C Hay, B. A. 2019. Cleave and Rescue, a novel selfish genetic element and general strategy for gene drive. Proc Natl Acad Sci, 116, 6250–6259.

Oberhofer, G., Ivy, T. C Hay, B. A. 2021. Split versions of Cleave and Rescue selfish genetic elements for measured self limiting gene drive. PLoS Genet, 17, e1009385.

Oberhofer, G., Johnson, M. L., Ivy, T., Antoshechkin, I. C Hay, B. A. 2024. Cleave and Rescue gamete killers create conditions for gene drive in plants. Nat Plants, 10, 936–953.

Raban, R., Marshall, J. M., Hay, B. A. C Akbari, O. S. 2023. Manipulating the Destiny of Wild Populations Using CRISPR. Annu Rev Genet, 57, 361–390.

Simoni, A., Hammond, A. M., Beaghton, A. K., Galizi, R., Taxiarchi, C., Kyrou, K., Meacci, D., Gribble, M., Morselli, G., Burt, A., Nolan, T. C Crisanti, A. 2020. A male-biased sex-distorter gene drive for the human malaria vector Anopheles gambiae Nat Biotechnol., 38, 1097–1097.

Smidler, A. L., Pai, J. J., Apte, R. A., Sánchez C. H. M., Corder, R. M., Jeffrey Gutiérrez, E., Thakre, N., Antoshechkin, I., Marshall, J. M. C Akbari, O. S. 2023. A confinable female-lethal population suppression system in the malaria vector, Anopheles gambiae. Sci Adv, 9, eade8903.

Sudweeks, J., Hollingsworth, B., Blondel, D. V., Campbell, K. J., Dhole, S., Eisemann, J. D., Edwards, O., Godwin, J., Howald, G. R., Oh, K. P., Piaggio, A. J., Prowse, T. A. A., Ross, J. V., Saah, J. R., Shiels, A. B., Thomas, P. Q., Threadgill, D. W., Vella, M. R., Gould, F. C Lloyd, A. L. 2019. Locally Fixed Alleles: A method to localize gene drive to island populations. Sci Rep, 9, 15821.

Thakkar, N., Hejzlarova, A., Brabec, V. C Dolezel, D. 2023. Germline Editing of Drosophila Using CRISPR-Cas9-Based Cytosine and Adenine Base Editors. The CRISPR Journal, 6, 557–569.

Tolosana, I., Willis, K., Burt, A., Gribble, M., Nolan, T., Crisanti, A. C Bernardini, F. 2025. A Y chromosome-linked genome editor for efficient population suppression in the malaria vector Anopheles gambiae. Nat Commun, 16, 206.

Vreysen, M. J., Saleh, K. M., Ali, M. Y., Abdulla, A. M., Zhu, Z. R., Juma, K. G., Dyck, V. A., Msangi, A. R., Mkonyi, P. A. C Feldmann, H. U. 2000. Glossina austeni (Diptera: Glossinidae) eradicated on the island of Unguja, Zanzibar, using the sterile insect technique. J Econ Entomol, 93, 123–35.

Willis, K. C Burt, A. 2021. Double drives and private alleles for localised population genetic control. PLoS Genet, 17.

Willis, K. C Burt, A. 2025. Engineering drive–selection balance for localized population suppression with neutral dynamics. Proc Natl Acad Sci, 122, e2414207122.

Wyss, J. H. 2000. Screwworm eradication in the Americas. Ann N Y Acad Sci, 916, 186–93.

Yang, E., Metzloff, M., Langmüller, A. M., Xu, X., Clark, A. G., Messer, P. W. C Champer, J. 2022. A homing suppression gene drive with multiplexed gRNAs maintains high drive conversion efficiency and avoids functional resistance alleles. G3: Genes Genomes Genetic, 12.

