## Supplementary Material for "Genomic editors for localised population suppression"

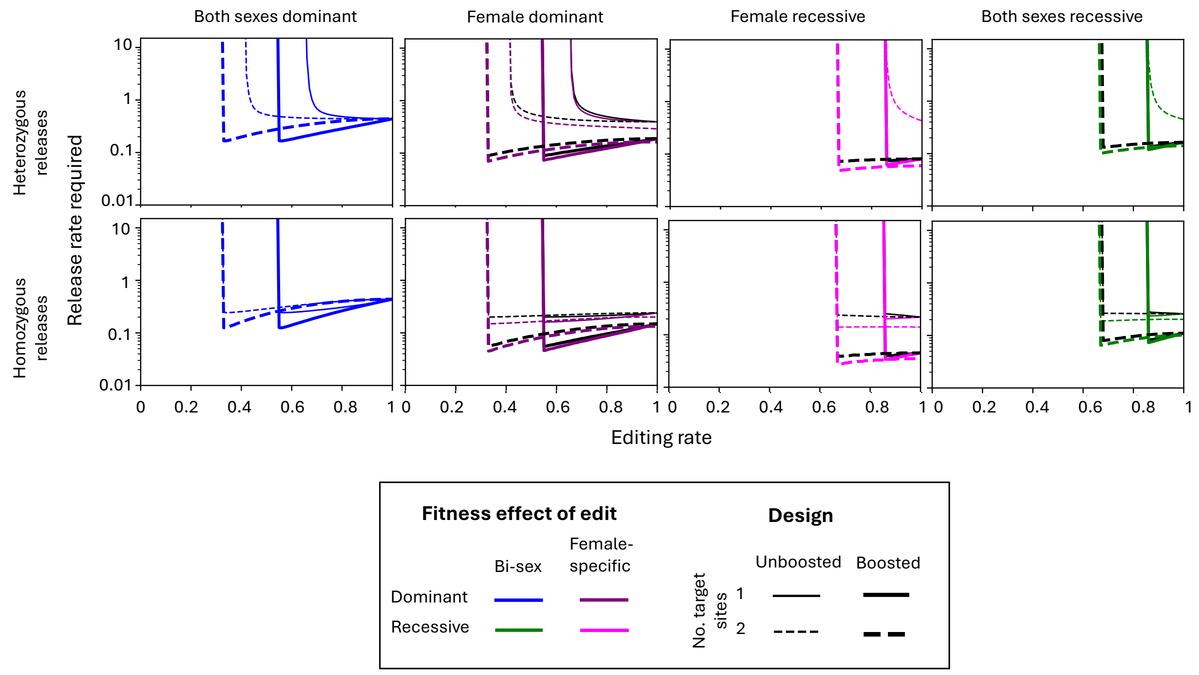


**Supplementary. Figure. 1** – As in Figure 3, the male release rates required relative to the starting male population size to suppress a population with an $R_m$ of 6 by 95% within 36 generations for a range of different autosomal editor strategies whilst varying the rate at which edits are created. Additional lines compared to figure 3 (dotted) and (dot-dashed) show designs where the edit target gene is linked to the editor, for one and two target sites respectively.
